# Mothers choice: maternal effects on the offspring population dynamics differed between exposure to microcystin-free algae *Microcystis aeruginosa* and chemical MC-LR environments in the rotifer *Brachionus calyciflorus*

**DOI:** 10.1101/2025.09.24.677296

**Authors:** Jia Na, Cuijuan Niu

## Abstract

Maternal effects can affect the offspring population dynamics by influence on the offspring’s phenotypes related to fitness, which is of great significance in regulation of population dynamics. The inhibitory effect of bloom algae Microcystis aeruginosa on the rotifer Brachionus calyciflorus has been extensively studied. However, weather algae M. aeruginosa or its toxin MC-LR will affect rotifer’s population dynamic via maternal effect is not clear yet. In this study, we investigated if fitness related phenotypes would change in offspring from mothers experienced microcystin-free M. aeruginosa or MC-LR stress environments by population dynamic, life history traits, and morphology respectively. We find that the survival and reproduction of offspring decreased under maternal microcystin-free M. aeruginosa stress, but increased under maternal MC-LR stress. Our results indicate that the maternal effects act on the offspring in different forms when faced with different environments. Our research further demonstrates that, unlike the common belief that algal toxins are the main factor inhibiting rotifer population growth, non-toxic Microcystis aeruginosa in fact shows a more potent suppressive effect.

## Introduction

Maternal effects occur when maternal phenotype and environment influence offspring phenotypic traits without altering their genotype (Hoyle & Ezard, 2012, Salinas, Brown & Mangel *et al*., 2013, Wenjie, Cuijuan & Xiaoxuan, 2019). As offspring traits—such as those related to survival and reproduction—directly shape fitness (Danchin, Charmantier & Champagne *et al*., 2011, Gilbert, Bosch & Ledón -Rettig, 2015),maternal effects play a critical role in shaping life history strategies and facilitating adaptive evolution of populations under natural selection pressures (Hoyle *et al*., 2012, Salinas *et al*., 2013, Wilson, Coltman & Pemberton *et al*., 2005).

Maternal effects are broadly divided into adaptive and non-adaptive types based on their impact on offspring fitness (Bernardo, 1996, Dustin J & Tobias, 2007). Adaptive maternal effects arise when mothers reliably anticipate the environmental conditions their offspring will encounter, thereby reducing mother–offspring conflict, generating offspring phenotypes that enhance fitness in the predicted environment (Dustin J *et al*., 2007) (Dustin J *et al*., 2007). For instance, *Daphnia magna* mothers exposed to microcystin-producing *Microcystis aeruginosa* produce offspring with elevated toxin tolerance (Radersma, Hegg & Noble *et al*., 2018). In contrast, non-adaptive maternal effects reduce offspring fitness as a result of diminished maternal investment in current reproduction (McCormick, 2006; Scheirs et al., 2000). Such effects typically occur under two circumstances: when mothers prioritize future reproductive opportunities over current ones, or when current environmental conditions threaten maternal survival, leading to resource allocation toward self-maintenance. In both cases, mothers sacrifice current reproductive output to maximize lifetime fitness (Dustin J *et al*., 2007). For example, *Pomacentrus amboinensis* female under high population density produce smaller offspring, which can subsequently impair population growth (McCormick, 2006). Similarly, parental exposure to endocrine-disrupting chemicals has been shown to disrupt offspring development, metabolism, and behavior, ultimately increasing mortality rates in the next generation (Schwindt, 2015).

Maternal effects can exert positive or negative influences on offspring, with their magnitude being strongly influenced by environmental conditions (Kuijper, Johnstone & Townley, 2014, Leimar & McNamara, 2015, Plaistow, Lapsley & Benton, 2006). This dependence is mediated by the accuracy of maternal predictions about future environments. Under stable conditions, where maternal and offspring environments are generally similar, these effects typically enhance offspring fitness. In rapidly changing environments, however, discrepancies between maternal predictions and actual offspring conditions often result in maladaptive phenotypes and reduced fitness (Kuijper *et al*., 2014, T, S & I, 2015). Marshall and Uller (2007) proposed that the adaptive value of maternal effects depends on several critical trade-offs: (1) maternal resource allocation versus survival, (2) environmental predictability versus variability, (3) mother-offspring conflicts in resource distribution and fitness interests, and (4) the cost-benefit balance associated with producing specific offspring phenotypes. Together, these interactions determine whether maternal effects ultimately improve or diminish offspring fitness (Dustin J *et al*., 2007) (Dustin J *et al*., 2007).

Population dynamics and their regulatory mechanisms represent fundamental questions in population ecology. Although classical approaches often assumed population homogeneity, modern ecological research emphasizes the importance of individual variation and its sources in understanding the regulation of population dynamics (Benton, St Clair & Plaistow, 2008, Moore, Whiteman & Martin, 2019, Stelzer & Snell, 2006). Maternal effects influence offspring phenotypes and life-history traits, particularly those associated with reproduction and survival, thereby contributing to variation in population dynamics (Love, McGowan & Sheriff, 2012, Yanagi & Tuda, 2010). For example, maternal size and age at reproduction affect both egg size and clutch size, which subsequently influence offspring survival and reproductive performance (Benton *et al*., 2008, García -Roger, Serra & Carmona, 2014).

Monogononta rotifer *B. calyciflorus* can rapidly expand its population and occupy ecological niches via parthenogenesis within a short time, making it an ideal model organism for studying the effects of maternal influences on population growth (Gilbert, 2004, Gilbert & Schröder, 2004). In natural water columns, algae serve as an essential food source for rotifers and can influence rotifer population dynamics through factors such as concentration, species composition, and nutritional quality. *Microcystis aeruginosa*, one of the most common cyanobacteria in algal blooms, occurs in two forms: microcystin-producing and microcystin-free.

The microcystin-producing *M. aeruginosa* generates cyclic heptapeptide hepatotoxins, known collectively as microcystins (MCs). Microcystins inhibit protein phosphatase activity. They are released into the water column following algal cell death and may be ingested by animals, exerting toxic effects on aquatic organisms, including zooplankton (III & Paerl, 1988, Liang, Gao & Shao *et al*., 2020, Liang, Ouyang & Chen *et al*., 2017, Soares, Lurling & Huszar, 2010). It is generally believed that *M. aeruginosa* influences rotifer populations through multiple interconnected mechanisms: (1) Microcystins (MCs): These toxins comprise several isomers, among which MC-LR is the most toxic and abundant during bloom events (Liang *et al*., 2020). (2) Nutrient composition. *M. aeruginosa* lacks critical unsaturated fatty acids essential for zooplankton growth (BRETT & MÜLLER-NAVARRA, 1997). (3) Particle size and morphology: Colonial forms of *Microcystis* are resistant to digestion and can physically interfere with rotifer filtration apparatus (de Bernardi & Giussani, 1990). (4) Inhibiting the growth of other algae: By releasing allelopathic compounds that suppress beneficial algae such as green algae—an important high-quality, *M. aeruginosa* indirectly impairs rotifer populations (Liang *et al*., 2017). (5) Other bioactive metabolites: Beyond microcystins, additional toxic compounds produced by *M. aeruginosa* adversely affect rotifers and other zooplankton (Jungmann, 1992, Rohrlack, Henning & Kohl, 1999, Starkweather & Kellar, 1987). Therefore, *M. aeruginosa* functions as a compound ecological factor with simultaneous and multifaceted effects. Taking *M. aeruginosa* as a single factor to explore its impact on zooplankton may not be able to reveal the underlying mechanism clearly.

This study aimed to examine how maternal effects influence offspring population dynamics in the *B*.*calyciflorus* when mothers are exposed to microcystin-free *M*.*aeruginosa* and MC-LR, respectively. Based on preliminary experiments, we predicted that the inhibitory effect of microcystin-free M. aeruginosa would be more pronounced than that induced by the pure chemical toxin MC-LR. Furthermore, we hypothesized that different environmental stresses experienced by mothers would lead to distinct maternal effects. We evaluated offspring population dynamics, life-table traits, and phenotypic parameters to assess fitness responses to maternal environmental stress in each treatment.

## Materials and methods

### 1. Experimental animals and culture conditions

The clonal strain of *B. calyciflorus* used in this study originated from a resting egg collected from Xihai Lake in Beijing, China (39°57′N,116°21′E). Clone has been continuously cultured in the laboratory under stable conditions, with resting eggs regularly collected. It has been genetically marked in a previous paper (mitochondria COI sequence in GenBank:MK344674 (Li *et al*. 2019)). Rotifers were cultured in COMBO medium (Kilham *et al*. 1998) at 20°C with a 16:8 (light: dark) photoperiod. During the acclimation phase, rotifers were fed with *C. Pyrenoidosa* at a concentration 1×10^6^ cells ml^-1^ and cultured under the above environments for at least a month before the experiment.

Two types of algae were utilized in the experiments: green algae *C. Pyrenoidosa* (daily culture food for rotifers) and blue algae Microcystin-free *M. aeruginosa* (experimental food in non-toxic cyanobacteria experiments), both cultured in BG-11 medium. Both the algae and medium recipes were sourced from the Institute of Hydrobiology (Chinese Academy of Science, Wuhan, China) and acclimated e in the laboratory at 25°C under a 16:8 (L:D) photoperiod.

### 2. Experimental chemicals

*Microcystin aeruginosa* MC-LR (Taiwan Algal Science Inc., ≥ 95% by HPLC) was stored as powdered solid in −20°C refrigerator, diluted to 1mg/mL with ultrapure water when being used, then further diluted to the desired concentration with COMBO medium in experiments.

### 3. Experimental design

Two experiments were conducted to explore the effects of maternal exposure to microcystin-free *M. aeruginosa* or MC-LR on population dynamics, life-history traits and phenotypic plasticity of the offspring.

**Experiment 1:** Effects of maternal exposure to microcystin-free *M. aeruginosa*

The 24h death rates of *B. calyciflorus* with the addition of varying concentrations of microcystin-free *M. aeruginosa* were used to determine LC50, which was found to be 8.861×10^4^cells ml^-1^. Consequently, we chose the level of 10^4^ cells ml^-1^ of microcystin-free *M. aeruginosa* (about 10% of the 24h LC50) as the treatment concentration. Details are shown in the supplemental materials. Specific experimental results are shown in Supplementary Material Fig. S2.

Newly born amictic females (F_0_) were picked and randomly assigned to COMBO medium at a density of 1 individual ml^-1^ with 0 (control, C) or 10^4^ cells ml^-1^ (blue algae exposure, M) microcystin-free *M. aeruginosa* added to the original food (1×10^6^cells ml^-1^ *C. Pyrenoidosa*). Offspring (F_1_) from each F_0_ group were collected immediately after birth and divided into four groups following the same operation as above: CC, CM, MC and MM (the first letter denotes the maternal treatment, and the second letter denotes the offspring treatment). Each of the four treatment groups, with six replicates per group, was then assessed for population dynamics, life table parameters and morphologic characters. For the experimental procedure flowchart, see Supplementary Material Fig. S1.

Population growth experiment was conducted in six-well plates. Young amictic females (<4h, F_1_) established populations at an initial density of 1 ind. ml^-1^ in one well with 8 ml experimental medium containing 1 × 10^6^ cells ml^-1^ *C. Pyrenoidosa* as food. Life table experiment was conducted in 24-well plates. Each well contained a single young amictic female (<4h, F_1_) with 1ml experimental medium. Six individuals constituted a cohort, with each experimental group having six replicate cohorts. Owing to experimental errors, only five replicates from the MC and MM groups were used for data analysis.

**Experiment 2:** Effects of maternal exposure to MC-LR

The 24h death rates of *B. calyciflorus* with the addition of various concentrations of MC-LR were used to calculate LC50, which was determined to be 265.410ng ml^-1^. The highest reported concentration of MC-LR in natural water bodies, 200 ng ml^-1^ (Lahti *et al*., 1997), was chosen as the treatment group concentration. Specific experimental results are shown in Supplementary Material Fig. S4.

Newly born amictic females (F_0_) were picked and randomly assigned to medium with 0 (control, C) or 200 ng ml^-1^ (MC-LR exposure, L) MC-LR added. Offspring (F_1_) from each F_0_ group were immediately collected after birth, then divided into four groups: CC, CL, LC and LL (the first letter denotes the maternal treatment, and the second letter denotes the offspring treatment). The present operations were the same as those described in experiment 1.

### 4. Measuring of population growth experiment

The total number of rotifers, including amictic females (AF) and mictic females (MF) were counted every 24h, until the population size stabilized. Female types were identified by the methods described in Snell and Carmona (1995). Population growth rate (r) was calculated with the formula *r* = (*ln*N_*t*_ − *ln*N_0_)/*t*, where N_0_ is the rotifer population density at the initial time (ind. ml^-1^), N_t_ is the rotifer population density at time point t (ind. ml^-1^), and t is the time taken for the population to reach its maximum (days). The mixis ratio (MR) was calculated by equation MR=MF/(MF^−^+AF).

### 5. Measuring of life table parameters

Rotifers were observed every 12h, recorded number of the survivor (F_1_) and the number of offspring produced (F_2_), then all numbered F_2_ were removed out from the experimental system. The medium was refreshed every 24 hours until all individuals of the F_1_ generation died. Life table parameters were calculated using following equations. Survival rate(l_x_) represents the percentage of surviving individuals, while age-specific reproductive rate(m_x_) represents the percentage of the offspring at age x(hours). Intrinsic growth rate (r_m_), representing the maximum instantaneous growth rate of the population under specific experimental conditions, was calculated using the gradual approximation method according to the equation 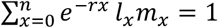. Total reproductive rate (G_0_) represents the number of offspring produced per female during the whole lifespan: *G*_0_ = ∑ *m*_*x*_. Net reproduction rate (R_0_) represents net growth rate of the cohort population through one generation, which was calculated by the equation *R*_0_ = ∑ *l*_*x*_ *m*_*x*_. Generation time (T) represents the average time to complete a generation, using the equation *T* = ln(R0)/*r*_*m*_ to get the exact value.

### 6. Measuring morphologic characters

Rotifers were cultured in 24-well plates. Each well contained a single young amictic female (<4h, F_1_) with 1ml experimental medium. Morphologic characters were measured at 48h (Most rotifers mature within 36h to 48h). Rotifers were fixed in a 5% formaldehyde solution, and their body length, body width and posterolateral spine length were measured under an electron microscopic imaging system (AxioCam MRc5 and Axioskop 2 plus, Carl Zeiss Inc., Germany). The body size (volume, V) of each rotifer was calculated by equation V = 0.52 × *a* ×*b* × *c*, where ‘a’ is body length, ‘b’ is body width, and ‘c’ is assumed to be 0.4a (Mccauley 1984).

### 7. Statistical analysis

Repeated measurement data, including offspring population density, mixis ratio, age-specific survival rate and age-specific reproduction rate, were analyzed using generalized estimating equations. Other data were tested for normality (Shapiro-Wilk test) and homogeneity of variance (Levene test), and those that met the quality for parametric test were analyzed using two-way ANOVA, while those did not meet were analyzed by Scheirer-Ray-Hare test. Significance level was set at p < 0.05. All statistical analyses were performed in R (R Core Team 2020, version 4.2.1).

To compare the extent of maternal effects on *B. calyciflorus* exposed to microcystin-free *M. aeruginosa* and MC-LR environments, Cohen’s d method was used to calculate and compare effect sizes for each. The influence of maternal effects on intrinsic growth rate and net reproduction rate under both environments was assessed by comparing the MM and CM groups, as well as the LL and CL groups, to determine the degree of difference. Calculations performed in R using the Cohen’s d function from the effsize package(Sullivan & Feinn 2012).

## Result

### 1 Effects of maternal exposure to microcystin-free *M. aeruginosa* on offspring population growth of *B. calyciflorus*

Population density was significantly affected by both maternal environment (F_0_) and offspring environment (F_1_), with a significant interaction (p<0.001;Table 1). However, the population growth rate (r) was only affected by maternal environment (p<0.001; 错误!未找到引用源。). Both population density and r values were depressed in all treatment groups with microcystin-free *M. aeruginosa* exposed history (CM, MC, MM; Fig 1A; Fig 1B). Rotifers whose mothers were not exposed to M (CC and CM) initially peaked on day 4. After passing the peak, population density of CC group declined and stabilized at a certain population size, while that of CM group decreased and crashed by day 10. Rotifers from mothers exposed to the M environment (MC and MM) showed a gradual decrease from the initial density (Fig 1A). Among the offspring exposed M environment, those from mothers with a M environment history did not vary much with those whose mothers exposed to the C environment (CM vs. MM, p=0.051; 错误!未找到引用源。 A). Population growth rate of the MM group was significantly lower than that of the CM group, even though both exhibited negative values (CM vs. MM, p=0.046;Fig 1B).

**Table 1:** Generalized estimating equation analysis of population, life table paraments for *B. calyciflorus* offspring under different treatments to microcystin-free *M. aeruginosa* exposure.

|  | Estimate | Std.err | Wald | p |
| --- | --- | --- | --- | --- |
| ● Population density |  |  |  |  |
| Maternal environment (A) | -2.687 | 0.192 | 195 | <0.001 |
| Offspring environment (B) | -2.310 | 0.197 | 137 | <0.001 |
| A×B | 2.385 | 0.207 | 132 | <0.001 |
| ● Mixis ratio |  |  |  |  |
| Maternal environment (A) | -0.199 | 0.056 | 12.53 | <0.001 |
| Offspring environment (B) | -0.035 | 0.066 | 0.27 | 0.601 |
| A×B | 0.074 | 0.088 | 0.70 | 0.402 |
| ● Survival rate |  |  |  |  |
| Maternal environment (A) | -0.189 | 0.068 | 7.67 | 0.006 |
| Offspring environment (B) | -0.123 | 0.061 | 4.04 | 0.044 |
| A×B | 0.209 | 0.095 | 4.87 | 0.027 |
| ● Age-specific reproductive rate |  |  |  |  |
| Maternal environment (A) | -0.244 | 0.070 | 12.13 | <0.001 |
| Offspring environment (B) | -0.146 | 0.075 | 3.77 | 0.052 |
| A×B | 0.186 | 0.091 | 4.17 | 0.041 |

**Fig 1:**
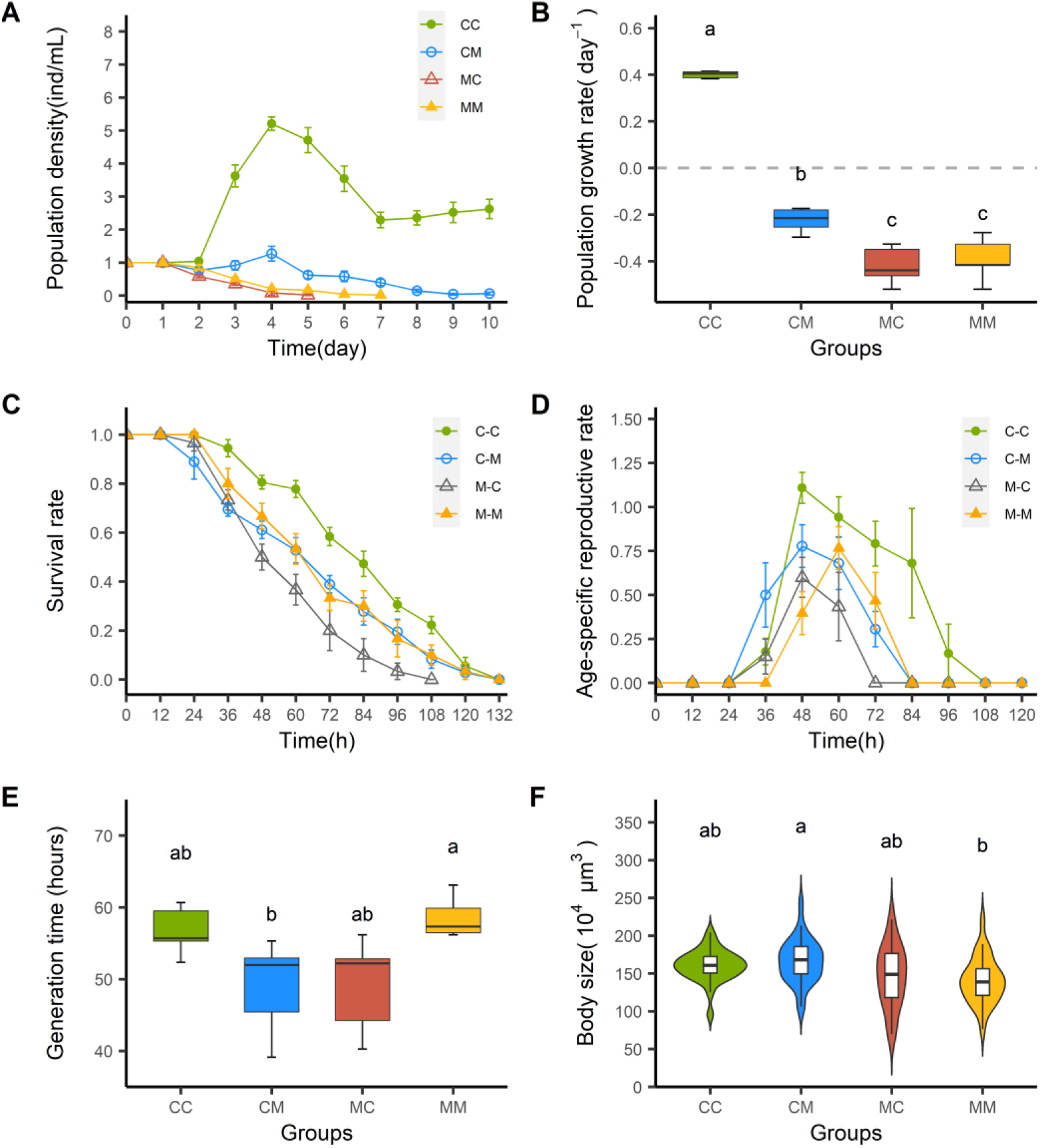
Population growth (A), population growth rates (B), survival rate (C), age-specific reproductive rate (D), generation time (E) and body size (F) of *B. calyciflorus* offspring under different treatments to microcystin-free *M. aeruginosa* exposure. The letter M means the addition of microcystin-free *M. aeruginosa*; C means non-addition of microcystin-free *M. aeruginosa*. In the combination of letters, the first letter represents environment experienced by mother, and the following letter represents environment experienced by offspring. Data are shown as mean ± SE. No common superscript letters represent significant difference between groups (P <0.05).

Maternal environment showed clear influence on the offspring’s life table and morphological parameters such as survival rate (p=0.006), age-specific reproductive rate (p<0.001), intrinsic growth rate (r_m_; p<0.001), total reproduction rate (G_0_; p=0.001), net reproduction rate (R_0_; p<0.001), life expectancy at hatching (e_x_; p<0.001), total offspring per female (p=0.003) and lifespan (p=0.024). Survival rate (p=0.027), age-specific reproductive rate (p=0.041), r_m_ (p<0.001), R_0_ (p=0.046), There were significant interactions between maternal and offspring environments on generation time (T; p<0.001), e^x^ (p<0.001), lifespan (p=0.007) and posterolateral spine length (p=0.024) of F_1_(Table 1;Table 2).

**Table 2:**
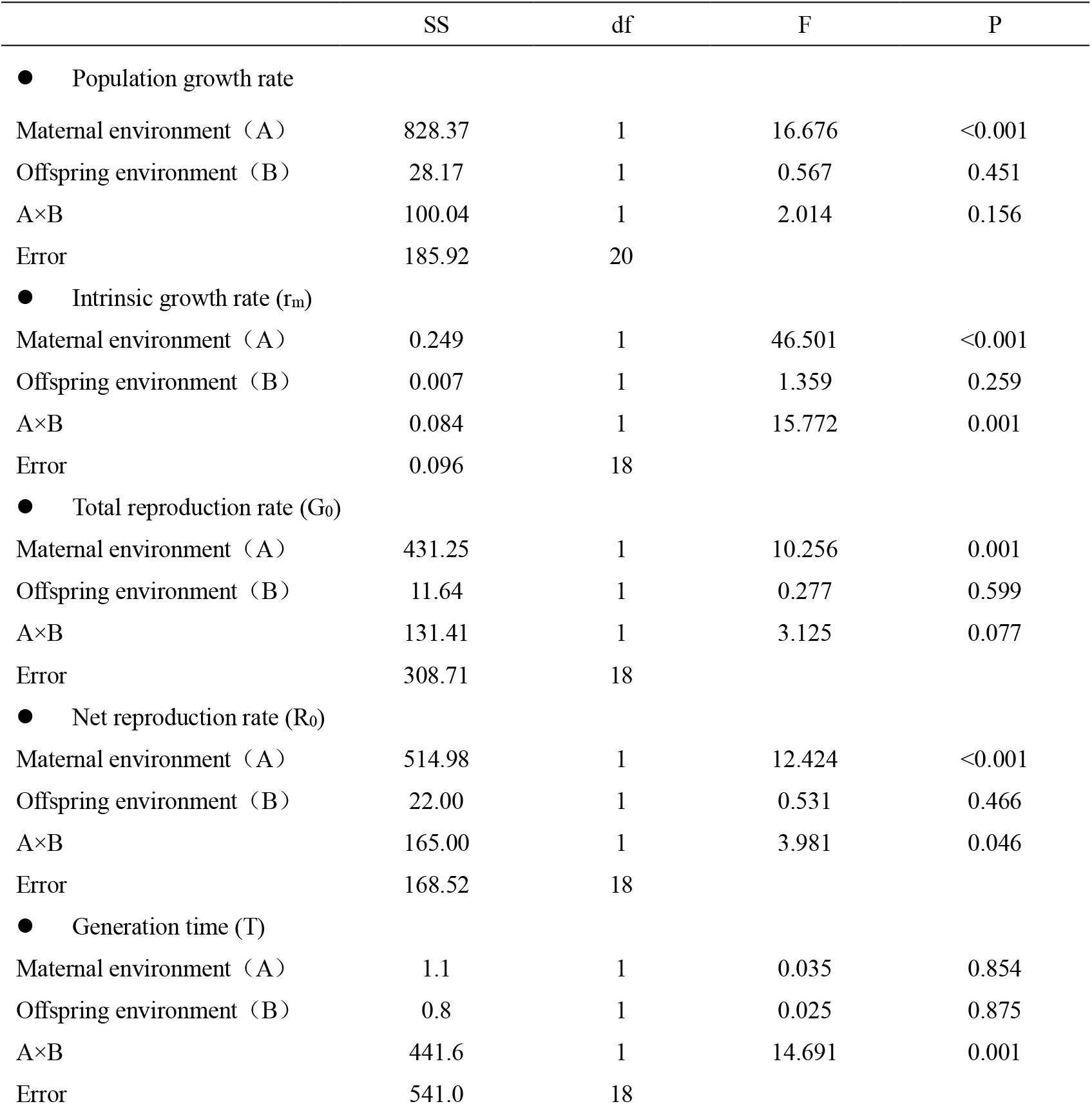
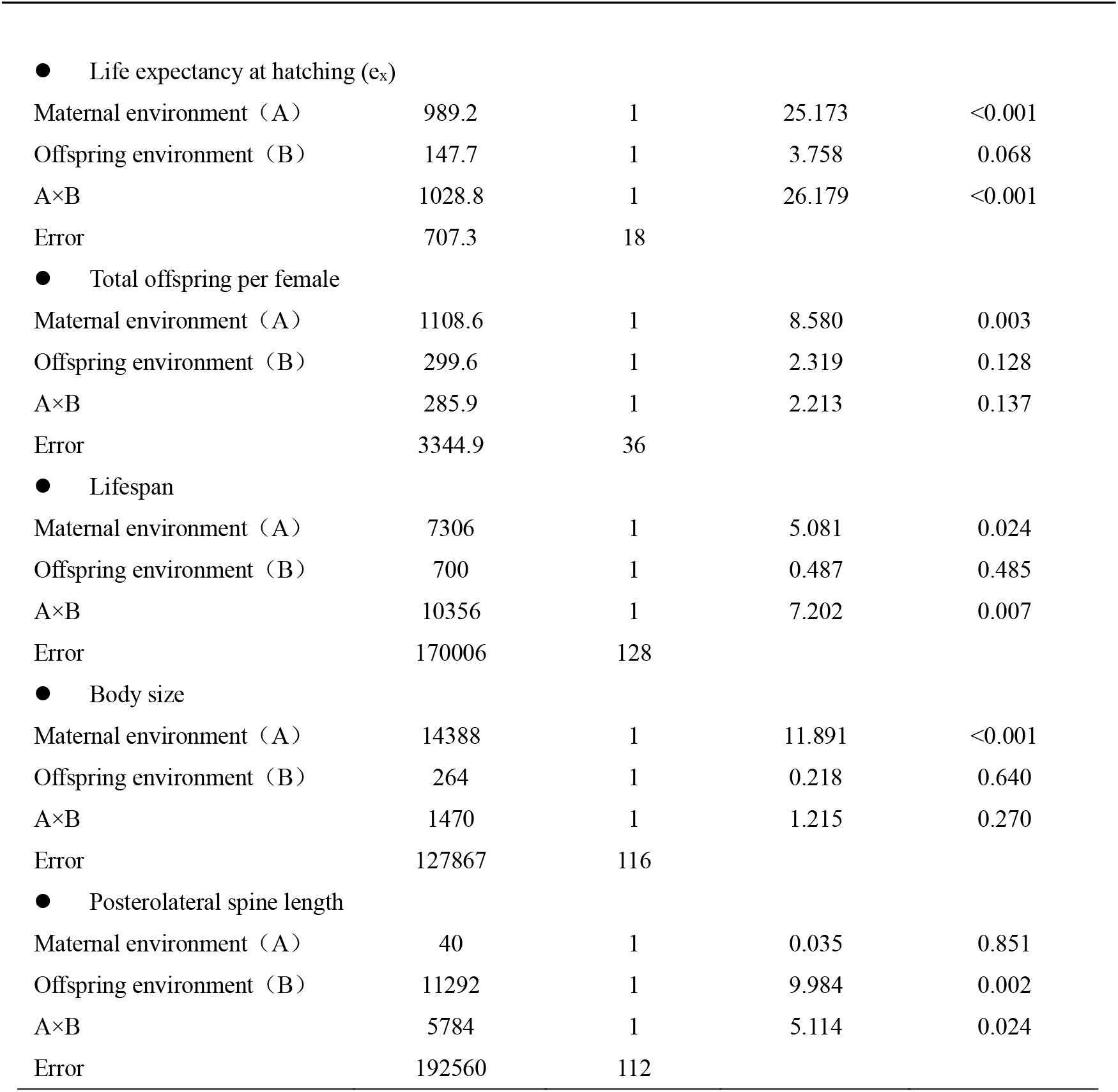
Two-way ANOVA of population, life table and morphological parameters for *B. calyciflorus* offspring under different treatments to microcystin-free *M. aeruginosa* exposure.

|  | SS | df | F | P |
| --- | --- | --- | --- | --- |
| ● Population growth rate |  |  |  |  |
| Maternal environment (A) | 828.37 | 1 | 16.676 | <0.001 |
| Offspring environment (B) | 28.17 | 1 | 0.567 | 0.451 |
| A×B | 100.04 | 1 | 2.014 | 0.156 |
| Error | 185.92 | 20 |  |  |
| ● Intrinsic growth rate ($r_m$) | | | | |
| Maternal environment (A) | 0.249 | 1 | 46.501 | <0.001 |
| Offspring environment (B) | 0.007 | 1 | 1.359 | 0.259 |
| A×B | 0.084 | 1 | 15.772 | 0.001 |
| Error | 0.096 | 18 |  |  |
| ● Total reproduction rate ($G_0$) | | | | |
| Maternal environment (A) | 431.25 | 1 | 10.256 | 0.001 |
| Offspring environment (B) | 11.64 | 1 | 0.277 | 0.599 |
| A×B | 131.41 | 1 | 3.125 | 0.077 |
| Error | 308.71 | 18 |  |  |
| ● Net reproduction rate ($R_0$) | | | | |
| Maternal environment (A) | 514.98 | 1 | 12.424 | <0.001 |
| Offspring environment (B) | 22.00 | 1 | 0.531 | 0.466 |
| A×B | 165.00 | 1 | 3.981 | 0.046 |
| Error | 168.52 | 18 |  |  |
| ● Generation time (T) |  |  |  |  |
| Maternal environment (A) | 1.1 | 1 | 0.035 | 0.854 |
| Offspring environment (B) | 0.8 | 1 | 0.025 | 0.875 |
| A×B | 441.6 | 1 | 14.691 | 0.001 |
| Error | 541.0 | 18 |  |  |
| ● Life expectancy at hatching ($e_x$) | | | | |
| Maternal environment (A) | 989.2 | 1 | 25.173 | <0.001 |
| Offspring environment (B) | 147.7 | 1 | 3.758 | 0.068 |
| A×B | 1028.8 | 1 | 26.179 | <0.001 |
| Error | 707.3 | 18 |  |  |
| ● Total offspring per female |  |  |  |  |
| Maternal environment (A) | 1108.6 | 1 | 8.580 | 0.003 |
| Offspring environment (B) | 299.6 | 1 | 2.319 | 0.128 |
| A×B | 285.9 | 1 | 2.213 | 0.137 |
| Error | 3344.9 | 36 |  |  |
| ● Lifespan |  |  |  |  |
| Maternal environment (A) | 7306 | 1 | 5.081 | 0.024 |
| Offspring environment (B) | 700 | 1 | 0.487 | 0.485 |
| A×B | 10356 | 1 | 7.202 | 0.007 |
| Error | 170006 | 128 |  |  |
| ● Body size |  |  |  |  |
| Maternal environment (A) | 14388 | 1 | 11.891 | <0.001 |
| Offspring environment (B) | 264 | 1 | 0.218 | 0.640 |
| A×B | 1470 | 1 | 1.215 | 0.270 |
| Error | 127867 | 116 |  |  |
| ● Posterolateral spine length |  |  |  |  |
| Maternal environment (A) | 40 | 1 | 0.035 | 0.851 |
| Offspring environment (B) | 11292 | 1 | 9.984 | 0.002 |
| A×B | 5784 | 1 | 5.114 | 0.024 |
| Error | 192560 | 112 |  |  |

Survival rate and age-specific reproductive rate were both the highest in CC group, but lowest in MC group among all four experimental groups,while those of CM and MM groups were similar (Fig 1C, D). Generation time of CM group was significantly shorter than that of the MM group (p=0.048; Fig 1E). Individual size of the CM group was significantly larger than that of the MM group (p=0.008; Fig 1F).

### 2 Effects of maternal exposure to MC-LR on offspring population of *B. calyciflorus*

Maternal and offspring environments showed significant influence and interaction on population density (p<0.001;Table 3) and population growth rate (r) (p<0.05;Table 4). CC group showed the maximum population density, peaking at day 5. Population growth of CL group was similar with that of LL group, whereas LC group showed the worst growth. LL and LC groups reached their maximum population densities most quickly, at day 2, while CL group peaked at day 3 (Fig 2A). The r value of LL group was significantly the highest among the experimental groups (LL vs other groups, p<0.05), while r value of LC group was clearly the lowest (LC vs. other groups, p<0.05; Fig 2B).

**Table 3:** Generalized estimating equation analysis of population, life table paraments for *B. calyciflorus* offspring under different treatments to MC-LR.

|  | Estimate | Std.err | Wald | p |
| --- | --- | --- | --- | --- |
| ● Population density |  |  |  |  |
| Maternal environment (A) | -2.796 | 0.286 | 95.8 | <0.001 |
| Offspring environment (B) | -2.112 | 0.305 | 47.9 | <0.001 |
| A×B | 2.819 | 0.362 | 60.5 | <0.001 |
| ● Mixis ratio |  |  |  |  |
| Maternal environment (A) | -0.0206 | 0.0430 | 0.23 | 0.63 |
| Offspring environment (B) | 0.0203 | 0.0402 | 0.25 | 0.61 |
| A×B | -0.0835 | 0.0600 | 1.93 | 0.16 |
| ● Survival rate |  |  |  |  |
| Maternal environment (A) | -0.093 | 0.070 | 1.76 | 0.18 |
| Offspring environment (B) | -0.056 | 0.065 | 0.74 | 0.39 |
| A×B | 0.113 | 0.096 | 1.40 | 0.24 |
| ● Age-specific reproductive rate |  |  |  |  |
| Maternal environment (A) | 0.119 | 0.080 | 2.21 | 0.14 |
| Offspring environment (B) | 0.029 | 0.067 | 0.18 | 0.67 |
| A×B | -0.006 | 0.120 | 0.00 | 0.96 |

**Table 4:**
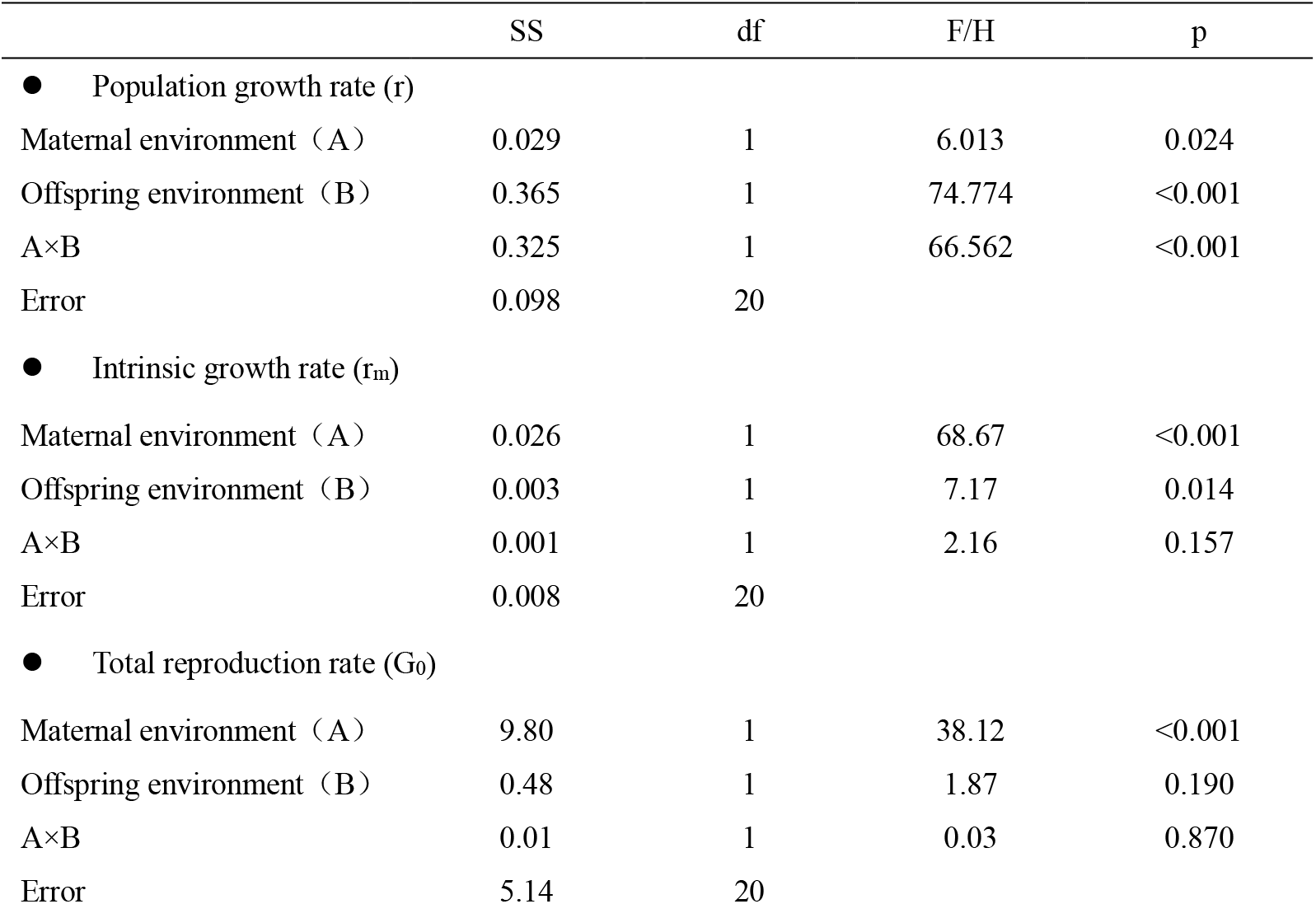
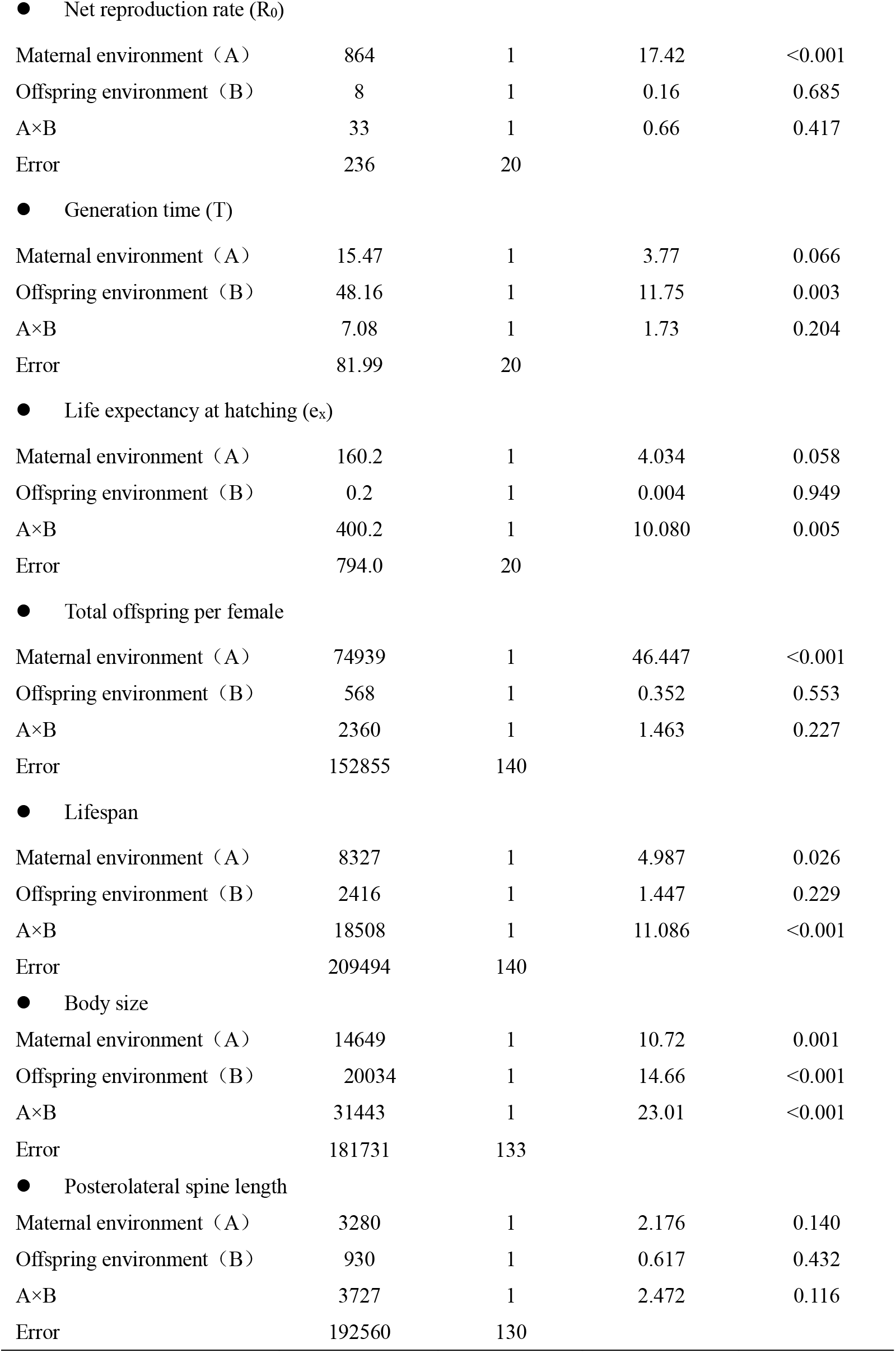
Two-way ANOVA of population, life table and morphological parameters for *B. calyciflorus* offspring under different treatments to MC-LR.

|  | SS | df | F/H | p |
| --- | --- | --- | --- | --- |
| ● Population growth rate (r) |  |  |  |  |
| Maternal environment (A) | 0.029 | 1 | 6.013 | 0.024 |
| Offspring environment (B) | 0.365 | 1 | 74.774 | <0.001 |
| A×B | 0.325 | 1 | 66.562 | <0.001 |
| Error | 0.098 | 20 |  |  |
| ● Intrinsic growth rate ($r_m$) | | | | |
| Maternal environment (A) | 0.026 | 1 | 68.67 | <0.001 |
| Offspring environment (B) | 0.003 | 1 | 7.17 | 0.014 |
| A×B | 0.001 | 1 | 2.16 | 0.157 |
| Error | 0.008 | 20 |  |  |
| ● Total reproduction rate ($G_0$) | | | | |
| Maternal environment (A) | 9.80 | 1 | 38.12 | <0.001 |
| Offspring environment (B) | 0.48 | 1 | 1.87 | 0.190 |
| A×B | 0.01 | 1 | 0.03 | 0.870 |
| Error | 5.14 | 20 |  |  |
| ● Net reproduction rate ($R_0$) | | | | |
| Maternal environment (A) | 864 | 1 | 17.42 | <0.001 |
| Offspring environment (B) | 8 | 1 | 0.16 | 0.685 |
| A×B | 33 | 1 | 0.66 | 0.417 |
| Error | 236 | 20 |  |  |
| ● Generation time (T) |  |  |  |  |
| Maternal environment (A) | 15.47 | 1 | 3.77 | 0.066 |
| Offspring environment (B) | 48.16 | 1 | 11.75 | 0.003 |
| A×B | 7.08 | 1 | 1.73 | 0.204 |
| Error | 81.99 | 20 |  |  |
| ● Life expectancy at hatching ($e_x$) | | | | |
| Maternal environment (A) | 160.2 | 1 | 4.034 | 0.058 |
| Offspring environment (B) | 0.2 | 1 | 0.004 | 0.949 |
| A×B | 400.2 | 1 | 10.080 | 0.005 |
| Error | 794.0 | 20 |  |  |
| ● Total offspring per female |  |  |  |  |
| Maternal environment (A) | 74939 | 1 | 46.447 | <0.001 |
| Offspring environment (B) | 568 | 1 | 0.352 | 0.553 |
| A×B | 2360 | 1 | 1.463 | 0.227 |
| Error | 152855 | 140 |  |  |
| ● Lifespan |  |  |  |  |
| Maternal environment (A) | 8327 | 1 | 4.987 | 0.026 |
| Offspring environment (B) | 2416 | 1 | 1.447 | 0.229 |
| A×B | 18508 | 1 | 11.086 | <0.001 |
| Error | 209494 | 140 |  |  |
| ● Body size |  |  |  |  |
| Maternal environment (A) | 14649 | 1 | 10.72 | 0.001 |
| Offspring environment (B) | 20034 | 1 | 14.66 | <0.001 |
| A×B | 31443 | 1 | 23.01 | <0.001 |
| Error | 181731 | 133 |  |  |
| ● Posterolateral spine length |  |  |  |  |
| Maternal environment (A) | 3280 | 1 | 2.176 | 0.140 |
| Offspring environment (B) | 930 | 1 | 0.617 | 0.432 |
| A×B | 3727 | 1 | 2.472 | 0.116 |
| Error | 192560 | 130 |  |  |

**Fig 2:**
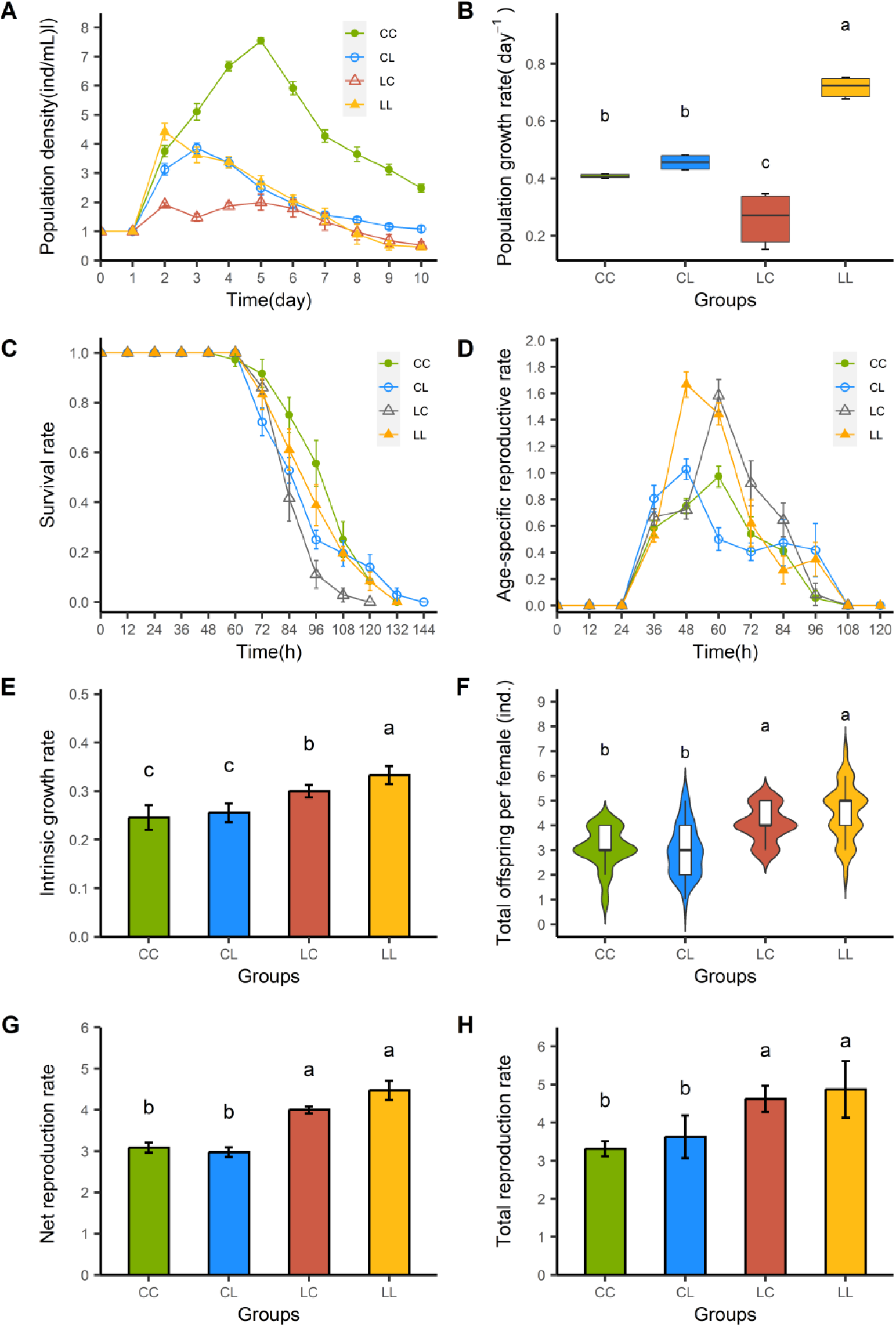
Population growth (A), population growth rates (B), survival rate (C), age-specific reproductive rate (D), intrinsic growth rate (E), total offspring per female (F), net reproduction rate (G) and total reproduction rate (H) of *B. calyciflorus* offspring under different treatments to microcystin-free *M. aeruginosa* exposure. The letter M means the addition of MC-LR; C means non-addition of microcystin-free *M. aeruginosa*. In the combination of letters, the first letter represents environment experienced by mother, and the following letter represents environment experienced by offspring. Data are shown as mean ± SE. No common superscript letters represent significant difference between groups (P <0.05).

In life table experiment, maternal environment significantly affects r_m_, R_0_, G_0_, total offspring per female (all p<0.001), and lifespan (p=0.026) of the offspring. Significant interactions between maternal and offspring environment were observed only for e_x_ (p=0.005) and lifespan (0<0.001;Table 4). CC group showed the best survival while LC group the worst, and that of LL group was similar to CL group (Fig 2C). Age specific reproduction rates of groups whose mothers exposed to L environment (LC and LL groups) were higher during 48-72h compared to those from mothers experienced C environment (CC and CL groups; Fig 2D). Neither maternal nor offspring environment showed significant effects or interaction on survival rate and age-specific reproductive rate curves under MC-LR treatment (Table 3). When mothers were not exposed to MC-LR, their offspring environment did not affect r_m_ of the population (CC vs. CL, p=0.827). However, when mothers were exposed to MC-LR, the offspring environment significantly affected the population’s r_m_ (LC vs. LL, p=0.038; Fig 2E). Regardless of the environment, offspring from L-exposed mothers showed significantly higher r_m_, R_0_, G_0_, and produced more offspring than those from C environment mothers (Fig 2E∼H). Compare to those in CL group, F_1_ in LL group had significantly higher population r_m_ (p<0.001), R_0_ (p=0.027), G_0_ (p=0.002), and produced more offspring F_2_ (p<0.001).

Body size of the offspring was significantly affected by both maternal and offspring environments, with a significant interaction (P<0.001;Table 4). Neither maternal nor offspring environments affected the posterolateral spine length, nor was there any interaction (p>0.05;Table 4). Among the treatment groups, body size of LC group was distinctly smaller than the other groups, and there was no significant difference between the others (supplementary table S2).

### 3 Maternal effect size under different treatment

For the r_m_ and R_0_, maternal exposure to microcystin-free *M. aeruginosa* exerts negative effects on offspring in the same stress environment. In contrast, maternal exposure to MC-LR environment positively influences offspring under similar stress conditions. The effect sizes of maternal exposure to both microcystin-free *M. aeruginosa* and MC-LR are highly significant for both intrinsic growth rate and net reproduction rate (p<0.001;Fig 3).

**Fig 3:**
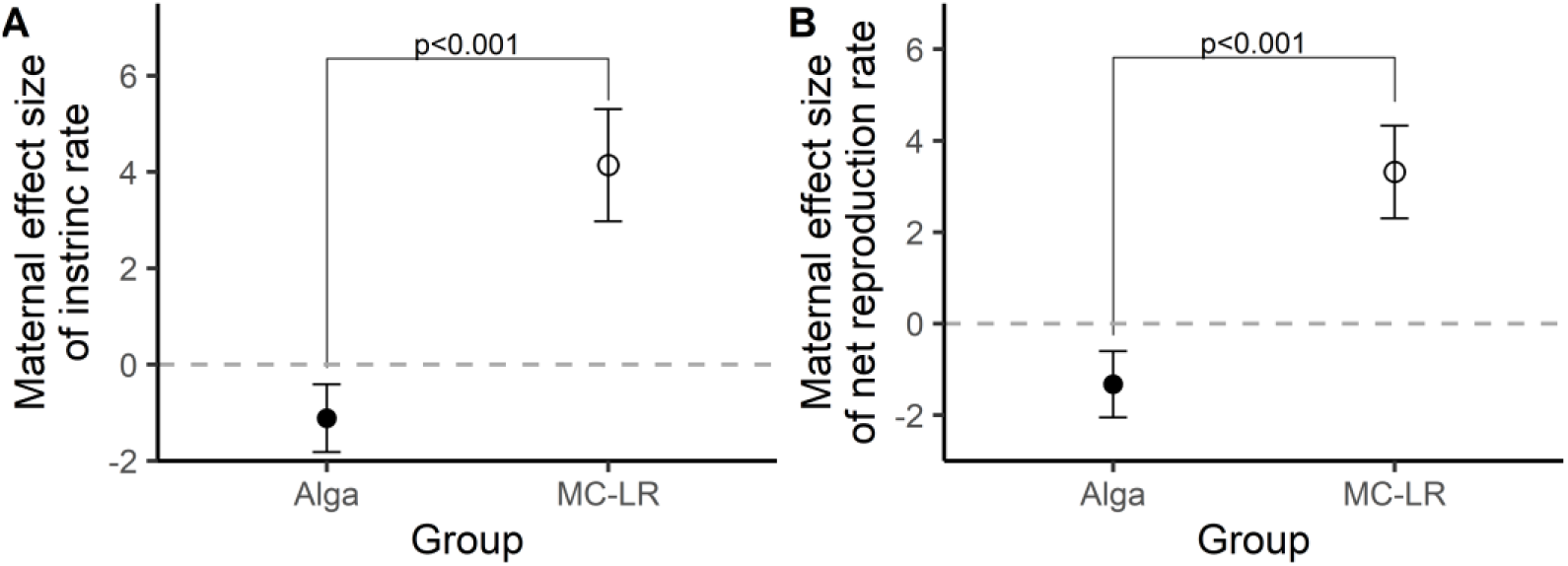
Maternal effect size of intrinsic growth rate (A) and net reproduction rate (B). ‘Alga’ represents the difference between MM group and CM group of *B. calyciflorus* offspring under microcystin-free *M. aeruginosa* exposure; ‘MC-LR’ represents the difference between LL group and CL group of *B. calyciflorus* offspring under MC-LR exposure. Data are shown as mean ± SE.

Fig 3 Maternal effect size of intrinsic growth rate (A) and net reproduction rate (B). ‘Alga’ represents the difference between MM group and CM group of *B. calyciflorus* offspring under microcystin-free *M. aeruginosa* exposure; ‘MC-LR’ represents the difference between LL group and CL group of *B. calyciflorus* offspring under MC-LR exposure. Data are shown as mean ± SE.

## Discussion

Contrary to the anticipatory maternal effect hypothesis, maternal exposure to microcystin-free *M. aeruginosa* did not enhance offspring fitness in the same environment. Compared with CM group, the MM group significantly grew slower, with a longer generation time and relatively lower fecundity (Fig 1E). Additionally, population density of MM group was consistently lower than that of CM group at the same time points (Fig 1A); body size of MM group was also significantly smaller than that of CM group (Fig 1F). Overall, the MM group demonstrated lower fitness in terms of both survival and reproduction compared to the CM group (supplementary table S1). These results suggest that maternal experience in a microcystin-free *M. aeruginosa* stress environment reduced offspring fitness in the same environment through selfish maternal effects.

Under harsh conditions, mothers face a trade-off between allocating resources to their own survival versus investing in reproduction. The ‘selfish maternal effect’ hypothesis suggests that when environmental conditions threaten maternal survival, mothers may prioritize self-maintenance, potentially at the expense of offspring fitness(Dustin J & Tobias 2007). Exposure to microcystin-free *M. aeruginosa* appears to elicit such a response, leading mothers to allocate energy to self-maintenance at the cost of offspring provisioning, thereby compromising offspring fitness. This interpretation is consistent with previous studies documenting maternal effects in rotifers fed exclusively with *M. aeruginosa* as food(Beyer & Hambright 2017; Hui *et al*. 2020). Our findings extend this understanding by demonstrating that rotifers exhibit non-adaptive maternal effects in response to *M. aeruginosa* even when the cyanobacterium produces no microcystins and adequate supplemental nutrition (*C. Pyrenoidosa*) is provided. Strikingly, even trace amounts of microcystin-free *M. aeruginosa* caused significant mortality that led to population collapse in *B. calyciflorus* offspring, despite the availability of sufficient suitable food.

Maternal exposure to microcystin-free M. aeruginosa significantly impaired key reproductive parameters, including G_0_, R_0_ and total offspring per female (Table 2). This reproductive decline likely resulted from suppressed survivorship. In the microcystin-free *M. aeruginosa* stress environment, rotifers matured at about 36 h and began producing offspring at about 36-48 h (Fig 1). However, survival rates for the three stress treatment groups (CM, MC, MM) were only 50-60% at 48 h (Fig 1C), implying that around half rotifers died during the pre-reproductive period, thus leading to decreased reproductive capacity.

Maternal exposure to MC-LR enhanced offspring fitness in the same environment, which supported the anticipated maternal effect hypothesis (Fig 2). This hypothesis proposes that offspring perform better when reared under conditions matching their maternal environment (Dustin J & Tobias 2007). The results confirmed that LL group showed a significantly higher r_m_, G_0_, R_0_ and total offspring per female compared to the CL group (Fig 2). No significant were observed in body size or posterolateral spine length between groups (supplementary table S2). Therefore, the LL group demonstrated higher fitness than the CL group in the MC-LR environment, lending support to the idea that maternal exposure can enhance offspring adaptation to a similar environment via anticipatory maternal effects. A key prerequisite for anticipatory maternal effects is that the maternal environment does not constitute a severe, life-threatening stressor. The MC-LR concentration used in this study meets this condition. Previous research has reported high tolerance in the rotifer *B. calycifloru*s to MC-LR, with concentrations below 200 ng mL^−1^ even potentially stimulating population growth (Huang *et al*. 2012). T This inherent tolerance likely ensured that maternal exposure under our experimental conditions was non-lethal, allowing mothers to invest resources in shaping offspring phenotypes, thereby potentially increasing offspring fitness.

This study demonstrates that various maternal effect responses occur under different environmental stressors (Fig 3). Mother faces a trade-off between its own fitness and that of offspring under stressful environments, responding with different maternal effect patterns based on the stress level. Under high stress levels, mother prioritizes its own survival over offspring fitness, leading to a ‘selfish maternal effect’; when the mother is tolerant to the environmental stressors, it enhances offspring fitness under the same stressful conditions through an ‘anticipated maternal effect’. In addition, we demonstrate that *M. aeruginosa* can inhibit *B. calycifloru*s growth even without producing microcystin, providing important evidence for future research.

## Conclusion

In conclusion, this study revealed contrasting transgenerational outcomes in *B. calyciflorus*: maternal exposure to microcystin-free *M. aeruginosa* environment reduced offspring fitness, whereas exposure to an MC-LR environment enhanced it. Specifically, Maternal microcystin-free *M. aeruginosa* environment led to decreased r, smaller body size, and prolonged T in offspring reared under the same condition. Furthermore, *B. calyciflorus* showed high sensitivity to microcystin-free *M. aeruginosa*, with even trace amounts markedly reduced survivability. In contrast, Maternal MC-LR environment exposure enhanced the offspring fitness in the same environment by increasing parameters related to growth, such as r_m_, and reproduction including G_0_, R_0_ and numbers of offspring. In summary, for M. aeruginosa, the alga itself and microcystin can affect B. calyciflorus offspring population dynamic via different type of maternal effects.

## Notes

### Competing Interest Statement

The authors have declared no competing interest.

### Funding

- National Natural Science Foundation of China: 32071528

