## supplementary for "Mothers choice: maternal effects on the offspring population dynamics differed between exposure to microcystin-free algae *Microcystis aeruginosa* and chemical MC-LR environments in the rotifer *Brachionus calyciflorus*"

Supplementary material

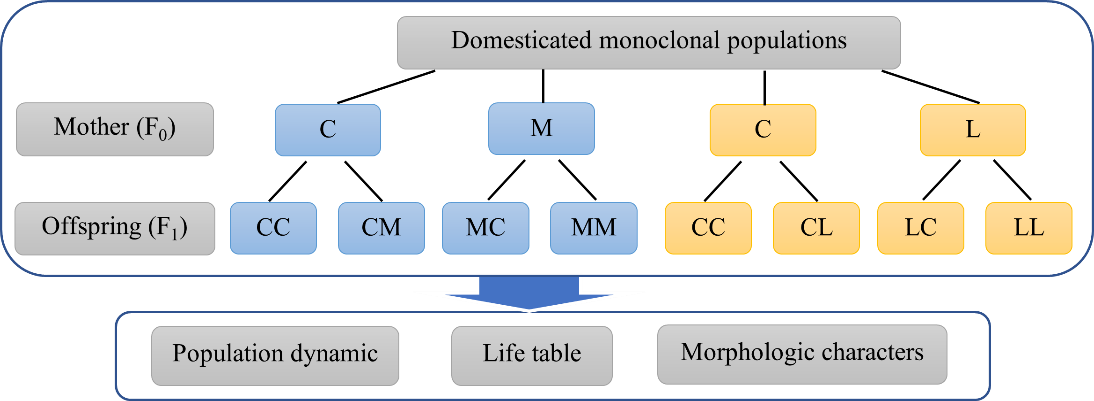

Fig S1 Experimental design. C represents the environment with *C. Pyrenoidosa*, M represents the environment with microcystin-free *Microcystis aeruginosa*, L represents the environment with MC-LR. The first letter of the two letter combinations represents the environment experienced by the mother (F_0_) and the second letter represents the environment experienced by the offspring (F_1_).

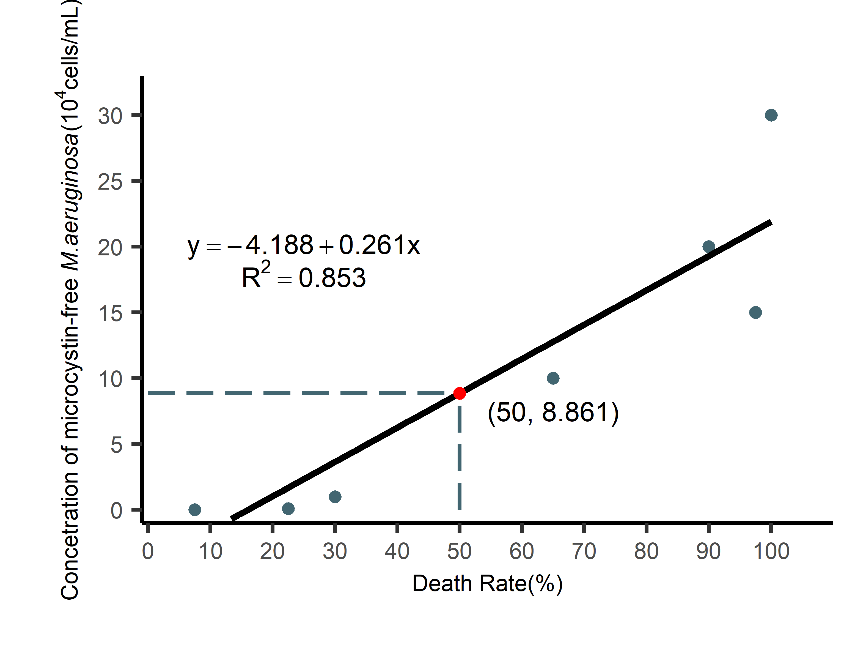

Fig S2 The 24h death rate of *B. calyciflorus* with addition of different concentrations of microcystin-free *M. aeruginosa*. The black dots represent the 24h lethal concentrations of rotifers at different concentrations, the solid black line represents the fitted straight line, the equation on the left is the fitted straight line equation and the coefficient of determination R^2^, and the red dot and coordinate are the LC50 calculated from the fitted straight line equation.

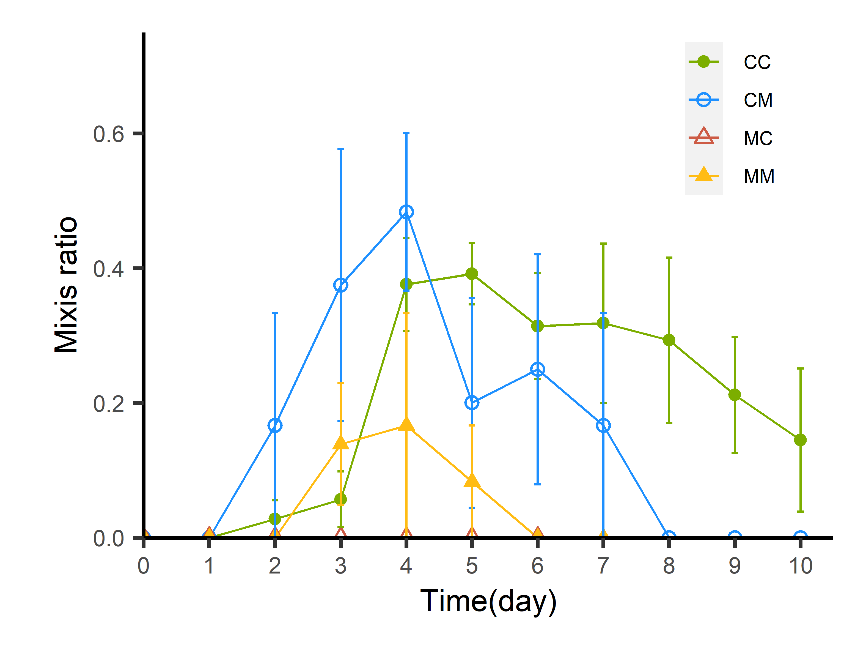

Fig S3 Changes of population mixis ratios of *B. calyciflorus* offspring under different treatments in the experiment of maternal effect of microcystin-free *M. aeruginosa*. Data points represent mean ± SE.

Table S1 Population, life history and morphological parameters of *B. calyciflorus* offspring under different treatments to microcystin-free *M. aeruginosa* exposure

| Treatment | | Population growth rate (Day^-1^) | Intrinsic growth rate (r_m_) | Total reproduction rate (G_0_) | Net reproduction rate (R_0_) | Generation time (T/h) |
| --- | --- | --- | --- | --- | --- | --- |
| Mother | Offspring |  |  |  |  |  |
| C | C | 0.400±0.014^a^ | 0.190±0.021^a^ | 3.867±0.540^a^ | 2.611±0.293^a^ | 57.717±1.898^ab^ |
|  | M | -0.222±0.019^b^ | 0.040±0.033^b^ | 2.264±0.266^ab^ | 1.250±0.129^b^ | 49.165±2.370^b^ |
| M | C | -0.437±0.029^c^ | -0.149±0.026^c^ | 1.183±0.138^b^ | 0.567±0.060^c^ | 49.159±2.650^ab^ |
|  | M | -0.385±0.040^c^ | -0.049±0.032^bc^ | 1.630±0.195^ab^ | 0.833±0.125^b^ | 58.602±1.160^a^ |
| Treatment | | Life expectancy at hatching (e_x_/h) | Total offspring per female (ind.) | Lifespan (h) | Body size  (10^4^μm^3^) | Posterolateral spine length (μm) |
| Mother | Offspring |  |  |  |  |  |
| C | C | 80.000±1.700^a^ | 5.530±0.403 | 84.333±4.840^a^ | 160.576±3.825^a^ | 63.125±3.591^ab^ |
|  | M | 62.333±3.142^b^ | 4.091±0.251 | 68.333±5.320^ab^ | 166.139±5.272^a^ | 66.342±4.197^ab^ |
| M | C | 52.800±2.440^b^ | 3.400±0.400 | 58.000±3.948^b^ | 146.229±7.340^ab^ | 51.787±4.095^b^ |
|  | M | 62.600±1.992^b^ | 3.571±0.202 | 70.400±5.358^ab^ | 140.801±5.568^b^ | 76.547±3.877^a^ |

Data are shown as mean ± SE. Different superscript letters represent significant differences between groups (p<0.05). C represents the environment without microcystin-free *M. aeruginosa*, M represents the environment with microcystin-free *M. aeruginosa*.

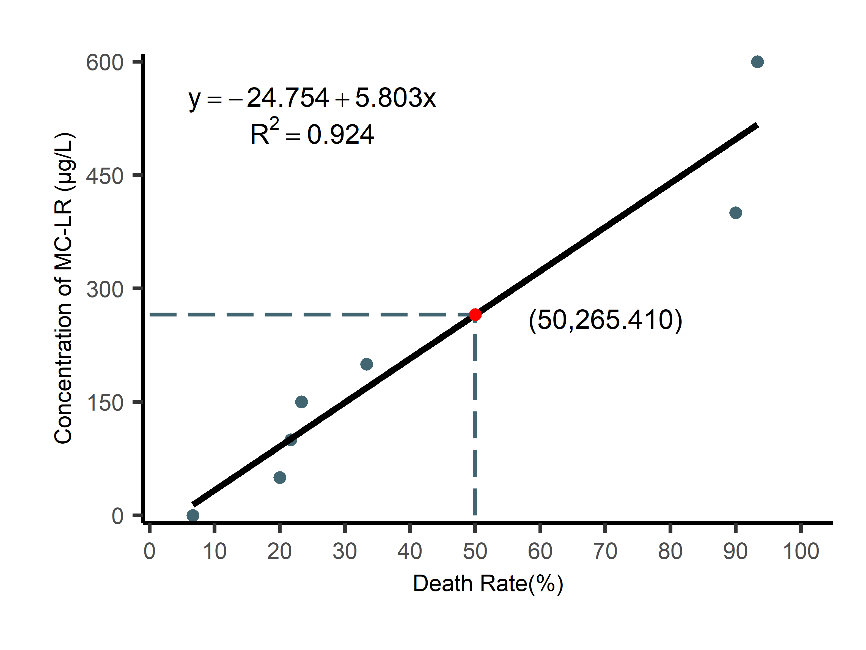

Fig S4 The 24h death rate of *B. calyciflorus* with addition of different concentrations of MC-LR. The black dots represent the 24h lethal concentrations of rotifers at different concentrations, the solid black line represents the fitted straight line, the equation on the left is the fitted straight-line equation and the coefficient of determination R^2^, and the red dot and coordinate are the LC50 calculated from the fitted straight line equation.

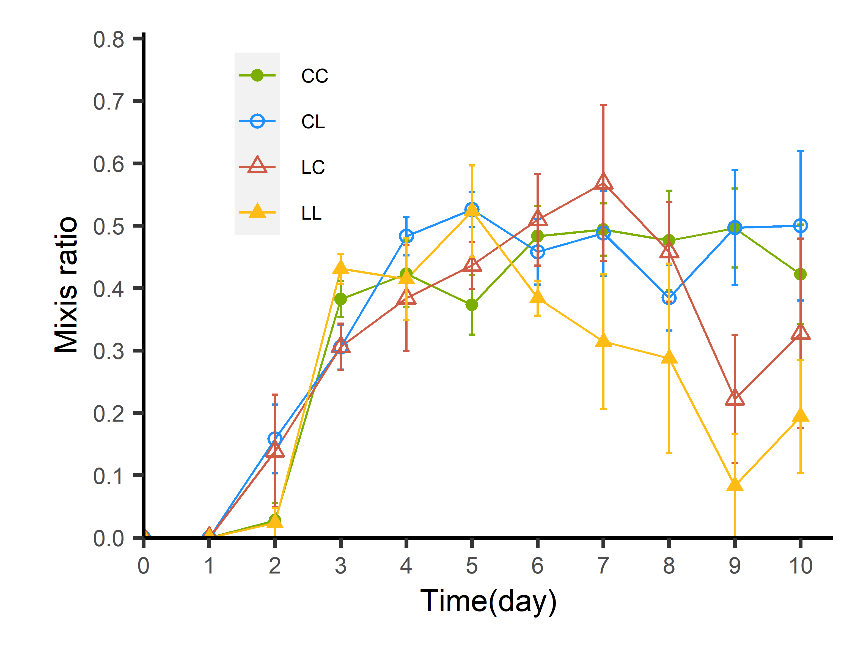

Fig S5 Changes of population mixis ratios of *B. calyciflorus* offspring under different treatments in the experiment of maternal effect of MC-LR. Data points represent mean ± SE.

Table S2 Population, life history and morphological parameters of *B. calyciflorus* offspring under different

treatments in the experiment of maternal effect of MC-LR.

| Treatment | | Population growth rate (Day^-1^) | Intrinsic growth rate (r_m_) | Total reproduction rate (G_0_) | Net reproduction rate (R_0_) | Generation time (T/h) |
| --- | --- | --- | --- | --- | --- | --- |
| Mother | Offspring |  |  |  |  |  |
| C | C | 0.421±0.016^b^ | 0.246±0.010^c^ | 3.311±0.074^b^ | 3.083±0.109^b^ | 54.930±0.664^a^ |
|  | L | 0.435±0.028^b^ | 0.255±0.007^c^ | 3.628±0.209^b^ | 2.972±0.107^b^ | 51.011±0.948^b^ |
| L | C | 0.258±0.037^c^ | 0.297±0.005^b^ | 4.620±0.154^a^ | 3.967±0.087^a^ | 55.630±0.586^a^ |
|  | L | 0.738±0.030^a^ | 0.334±0.008^a^ | 5.047±0.274^a^ | 4.567±0.234^a^ | 54.298±0.739^ab^ |
| Treatment | | Life expectancy at hatching (e_x_/h) | Total offspring per female (ind.) | Lifespan (h) | Body size  (10^4^μm^3^) | Posterolateral spine length (μm) |
| Mother | Offspring |  |  |  |  |  |
| C | C | 96.333±3.445^a^ | 3.083±0.134^b^ | 94.333±2.669^a^ | 213.954±5.799^a^ | 93.243±5.193 |
|  | L | 88.333±1.661^ab^ | 2.972±0.167^b^ | 82.333±3.518^b^ | 208.524±5.061^a^ | 79.739±5.702 |
| L | C | 83.200±2.504^b^ | 4.056±0.119^a^ | 77.000±1.875^b^ | 161.710±5.634^b^ | 92.434±5.601 |
|  | L | 92.000±1.960^ab^ | 4.500±0.185^a^ | 85.333±3.053^b^ | 216.924±8.186^b^ | 96.705±4.122 |

Data are shown as mean ± SE. Different superscript letters represent significant differences between groups (p<0.05). C represents the environment without MC-LR, L represents the environment with MC-LR.
